# e-workflow for recording of glycomic mass spectrometric data in compliance with reporting guidelines

**DOI:** 10.1101/401141

**Authors:** Miguel A. Rojas-Macias, Julien Mariethoz, Peter Andersson, Chunsheng Jin, Vignesh Venkatakrishnan, Nobuyuki P. Aoki, Daisuke Shinmachi, Christopher Ashwood, Katarina Madunic, Tao Zhang, Rebecca L. Miller, Oliver Horlacher, Weston B. Struwe, Fredrik Levander, Daniel Kolarich, Pauline M. Rudd, Manfred Wuhrer, Carsten Kettner, Nicolle H. Packer, Kiyoko F. Aoki-Kinoshita, Frédérique Lisacek, Niclas G. Karlsson

## Abstract

Glycomics targets released glycans from proteins, lipids and proteoglycans. High throughput glycomics is based on mass spectrometry (MS) that increasingly depends on exchange of data with databases and the use of software. This requires an agreed format for accurately recording of experiments, developing consistent storage modules and granting public access to glycomic MS data. The introduction of the MIRAGE (Mimimum Requirement for A Glycomics Experiment) reporting standards for glycomics was the first step towards automating glycomic data recording. This report describes a glycomic e-infrastructure utilizing a well established glycomics recording format (GlycoWorkbench), and a dedicated web tool for submitting MIRAGE-compatible MS information into a public experimental repository, UniCarb-DR. The submission of data to UniCarb-DR should be a part of the submission process for publications with glycomics MS^n^ that conform to the MIRAGE guidelines. The structure of this pipeline allows submission of most MS workflows used in glycomics.

## INTRODUCTION

Posttranslational modifications of proteins play an essential role in modifying the single 20 amino acids used in protein synthesis thereby extending the functions and regulating the activities of the synthesized proteins. A census of all possible protein forms now commonly called proteoforms was recently estimated^1^. In this renewed view of protein diversity, glycoforms are expected to play a major role. Glycosylation is the basis of several biological events that include structural stability, recognition, immunological responses, cancer development and the attachment of pathogens to host cells as the first step in the process of infection^2, 3^. Furthermore, defects in glycosylation result in a range of human diseases, including Congenital Disorders of Glycosylation (CDG) caused by mild-loss of function in *N*-linked^4^ and *O*-linked oligosaccharide^5^ biosynthesis, both of whish result in sever illness, organ failure and premature death.

Mass spectrometry (MS) has become the central tool for the study of protein glycosylation largely due to its speed, high sensitivity and the structural information it delivers^6, 7^. Due to the range in experimental techniques and the nature of MS systems there are therefore large amount of data generated. Glycan profiling by MS of free and released glycans makes use of a considerable variety of experimental techniques and instrumentation to increase speed, depth and efficiency. This is summarized in Figure 1 (adapted from^8^), where a generic workflow for glycan analysis is shown as well as the different options available for each task.

**Figure 1.**
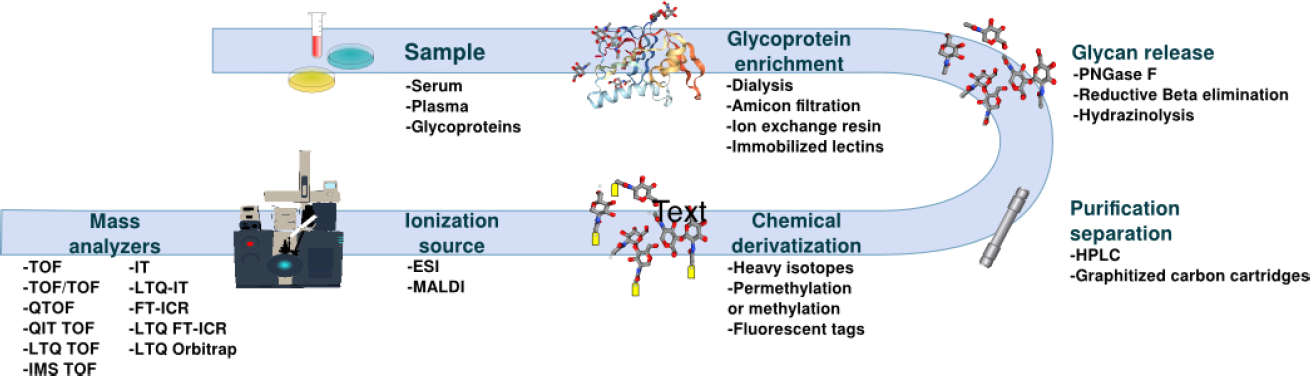
General Overview of a workflow for a glycomic MS workflow.

A factual assessment process to acknowledge the existence of a literature reported glycan and its localization (e. g. tissue and/or protein) demands a thorough description of experimental conditions and instrumental settings. The reproducibility of results also requires a disclosure of how the structural assignment was performed, e.g., manually or software assisted. Therefore, to ensure transparency and reproducibility there is a need for comprehensive reporting of experimental methods in publications. Other omics fields have addressed these concerns, which motivated the respective science communities to develop guidelines for the reporting, collecting and distributing of data and information. This started with MIAME launched for the handling of microarrays^9^. Shortly after was the arrival of MIAPE guidelines for proteomics^10^. Since these early efforts, many guidelines were published in a wide range of applications such as STRENDA in enzymology^11^ or CIMR in metabolomics^12, 13^. There are currently 120 reporting guidelines published and registered with the BioSharing portal^14^. The glycomics community has proposed the MIRAGE (Minimum Information Required for A Glycomics Experiment) project in 2011 supported by experts from the diverse areas of glycomics research^15^. This has resulted in setting up the following set of guidelines: sample preparation^16^(doi:10.3762/mirage.1), MS analysis^17^ (doi:10.3762/mirage.2), glycan microarray analysis^18^ doi:10.3762/mirage.3), and liquid chromatography guidelines (manuscript in preparation).

These glycomic guidelines need to be viewed in a larger picture of e(lectronic)-infrastructure. In 2008 an NIH “Frontiers in glycomics” work group stressed in its white paper the need for a curated glyco-structure database with long term funding^19^. This database should contain associated information about experimental and biosynthetic data. Attempts to create this “super” glycomic database was pioneered already in the 80s with Carbank^20^, and has been followed by GlycomeDB^21^, GLYCOSCIENCES.de^22^, GlycosuiteDB^23^ and UniCarbKB^24^. The latest release of GlyConnect (https://glyconnect.expasy.org) at the Swiss Institute for Bioinformatics provides a long-term solution for a stable and financially supported database of glycoconjugates (manuscript in preparation). There is also now an agreement in the glyco-research field, that all proposed glycan structures should be registered with a unique identifier in the GlyTouCan registry. This provides a foundation for developing complementary repositories, where unique glycan records can be associated with additional information, including incorporation of the MIRAGE reporting guidelines.

Presently, glycomic experimental MS data are collected and stored locally in individual research laboratories. Data sharing between different labs that may perform comparable experiments yet subtle differences in methods often precludes distribution because of incomplete recording of experimental protocols as well as conflicting hardware and software parameters. However, glycomics is practiced in laboratories worldwide that address glycobiological questions in cancer, inflammation and infection to name only a few and critically they require tools to share their findings and data. Especially now, when powerful software tools can interpret large complex MS datasets, making it impossible disclose all data as part of a traditional publication. The database UniCarb-DB was set up in 2009^25, 26^ in order to set up the framework for providing access to experimental MS data including both fragmentation spectra with associated structures and metadata about biological origin. Currently, it contains structural and fragmentation data of O-glycans, N-glycans and glycosaminoglycans obtained in positive and negative ion modes. Since its introduction, several versions of UniCarb-DB have been released, mainly to improve the glycomic data quality, increase the number of entries and advance the usability of the application. Reference MS fragmentation spectra of glycans are also being assembled at the NIST Glycan Mass Spectral Reference Library (https://chemdata.nist.gov/glycan/spectra).

The progress in glycomic software development for glycomics has been slow but steady. The early GlycosidIQ automated the comparison of observed fragments with theoretical glycofragments derived from a structural database^27^. This approach has been adopted in commercial software^28^, and expanded into matching glycopeptide data in the Glycomaster DB application^29^. Other approaches convert spectra into structures relying on spectral libraries^26, 30^. More advanced tools for glycomics use partial *de novo* sequencing^31^ including the recently published Glycoforest^32^. High throughput glycomic annotation (GRITS Toolbox, www.grits-toolbox.org/) and quantitation tools^33^ in glycomics are now available and increase the need for a common data exchange format, for these data to be publicly accessible and scrutinized. If these data can be provided in an agreed format, the validation and curation of structures deposited into the GlyTouCan registry will become feasible.

In this paper, we propose a workflow to collect, process and store experimental data in compliance with the MIRAGE MS and sample preparation guidelines a UniCarb-DR (DR = Data Repository) that benefits from the previous developed UniCarb-DB framework of quality LC-MS/MS data and structural assignments^25, 26^. UniCarb-DR incorporates both the MIRAGE MS and sample preparations guidelines. It also provides an electronic submission tool, guiding users for initial data validation to ensure all required information is provided. Data is entered in a structured form (template, http://unicarb-dr.biomedicine.gu.se/generate) that can be submitted to UniCarb-DR together with GlycoWorkbench files, including structures, spectra, fragmentation annotation and meta-data with scoring parameters, spectral quality and the use of orthogonal methods for structural assignments.

## METHODS

### System overview and implementation

UniCarb-DR repository is based on the UniCarb-DB database format^25, 26^, adopted to include tables and layouts for MIRAGE information. The repository design is based on a PostgreSQL as database manager system. The UniCarb-DR web application is supported by the Play Framework (**<u>Error! Hyperlink reference not valid.</u>** The Play Framework makes use of the MVC paradigm, where the elements of an application adopt one of three roles: Model, View or Controller. The Model is written in Java and represents the data and how the data is manipulated. The View is the layer that is displayed to users in the web interface. In UniCarb-DR, the View is written in Scala, JavaScript and implements the Jquery, Bootstrap and SpeckTackle libraries for data visualization. The Controller layer, also written in Java, controls the data that flows to the model and updates the View when the data change in response to user actions.

### Testing of the MIRAGE glycomic workflow

In order to develop and test the MIRAGE parameter on-line form and the submission tool, we selected beta-test sites that generated glycomic LC-MS^2^ and MS^2^ from *N*-linked, *O*-linked and proteoglycan type protein oligosaccharides ((http://unicarb-dr.biomedicine.gu.se/references). MIRAGE data spreadsheets were generated via the described on-line submission form available at http://unicarb-dr.biomedicine.gu.se/generate, where LC parameters also were recorded. Generated spreadsheets from this submission are available in supplementary material. Individual centroided MS^2^ spectra were copied manually into GlycoWorkbench^34^.gwp files together with the identified structures assigned from peak matching or manual interpretation Examples of Glycoworkbench files is also available in Supplementary material. Structures were assigned based on MS^2^ spectra and/or retention time and the quality of matching was manually validated. Reference LC-MS .raw were uploaded to Swegrid via the Proteios submission portal, also converting the data into .mzml format.

## RESULTS

### UniCarb-DR application to support using MIRAGE guidelines

In order to implement the glycomic MIRAGE guidelines^15^, we identified and adopted the two existing guidelines for glycomic sample preparation^16^ and MS^17^. In addition, an HPLC experimental recording was developed, expanding on the guidelines to enable recording of LC-MS parameters. For this first version, efforts targeted to the most essential implementations (the qualitative/structural information) and thus the quantification aspects of the guidelines were only addressed on a superficial level. This is justified considering that workflows for quantitative glycomics are still evolving and the basic level of methods and software tools are unavailable.

To support the e-workflow, we created the UniCarb-DR (Data Repository) (http://unicarb-dr.biomedicine.gu.se/) that facilitates submission of glycomics MS^n^ data conforming to the MIRAGE guidelines as part of a publication submission process (Figure 2). This will serve as the interim storage of experimental MS fragment data and structures before curation and transition into the UniCarb-DB database. A submission tool allows the user to browse and re-enter submitted data before it is uploaded to the repository. Since most data is likely to be submitted before publication, the user will need to refer to it as a “manuscript”. If data is uploaded after publication, the PubMed ID (PMID) available from (https://www.ncbi.nlm.nih.gov/pubmed/) will be required. The deposition of data in the repository first requires the registration and login as a user at http://unicarb-dr.biomedicine.gu.se/login and then providing a number of files and information (Figure 2) including:

1. A compiled file with MIRAGE data (proposed format is spreadsheets).
2. Compiled information about structures (proposed format is GlycoWorkbench)^34^.
3. Location of publicly accessible unprocessed MS files
4. Unique structure identifier (this information is automatically generated by communication with GlyTouCan structural repository^35^).

**Figure 2.**
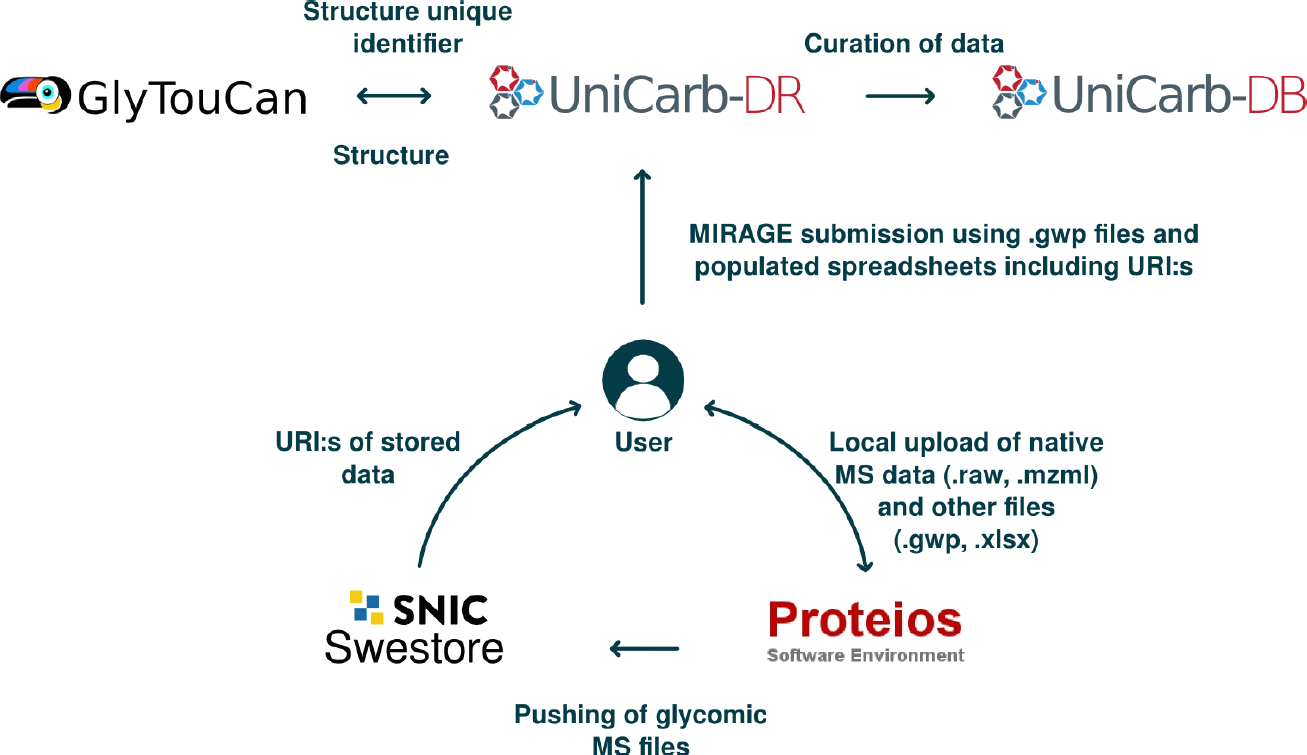
Overview of the workflow for MIRAGE data preparation and submission to UniCarb-DR. The workflow was designed to support the MIRAGE guidelines for sample preparation MS, as well as HPLC information. General features that are relevant to the entire experiment can be captured by means of automatically generated spreadsheets and/or online forms, accessible on the UniCarb-DR website. GlycoWorkBench files are proposed to be used for data on individual structures. Native glycomics MS data in this report was uploaded via Proteios and converted into mzML format. These together with MIRAGE files were pushed to Swestore, that generated publically available URI:s for individual files, as an example for how to make native data publically available These URI:s can be provided in the MIRAGE spreadsheets. A submission tool is available from UniCarb-DR, where the collected data are evaluated, stored and posted on the website with restricted access. UniCarb-DR also communicates with the global glycostructure repository GlyTouCan to generate unique identifiers for structures submitted to UniCarb-DR. UniCarb-DR will become one curation source of data for UniCarb-DB.

The specific information for each of these four items is described below.

### Step 1) MIRAGE record-Spreadsheets

Experimental data needs to be provided in a spreadsheet form with data fields reflecting the principle structure of the MIRAGE guidelines. Prefilling and downloading of the MIRAGE compliant spreadsheets is possible in the web form (http://unicarb-dr.biomedicine.gu.se/generate). Three different spreadsheets are available: 1) sample preparation, 2) LC and 3) MS guidelines. These can be generated individually or combined into one file with several sheets. These spreadsheets can be modified off-line using common software packages such as Excel template v 2007 or later (saved as xlsx file type).

### Step 2) MIRAGE record-GlycoWorkbench

The open source software GlycoWorkbench developed by EuroCarbDB to assist manual interpretation of MS data^34^ is used to generate initial glycan structures. Glycoworkbench provides a straightforward interface to draw glycan structures in different cartoon formats (SNFG, Oxford, IUPAC) using the embedded GlycanBuilder module^34^. Glycan structures are stored in a linear format (.gws) for easy parsing and recording into databases. All recorded data is stored in an XML format (.gwp file extension) (Figure 3). GlycoWorkbench allows the recording of individual structures as a “Scan” with associated fragment data (fragment list imported from MS software as centroided data). The first level should be MS^2^. Hence, a GlycoWorkbench file that includes both several structures and MS^2^ data needs to be defined by several “Scans” (one for each structure) directly under the “Workspace” item.

**Figure 3.**
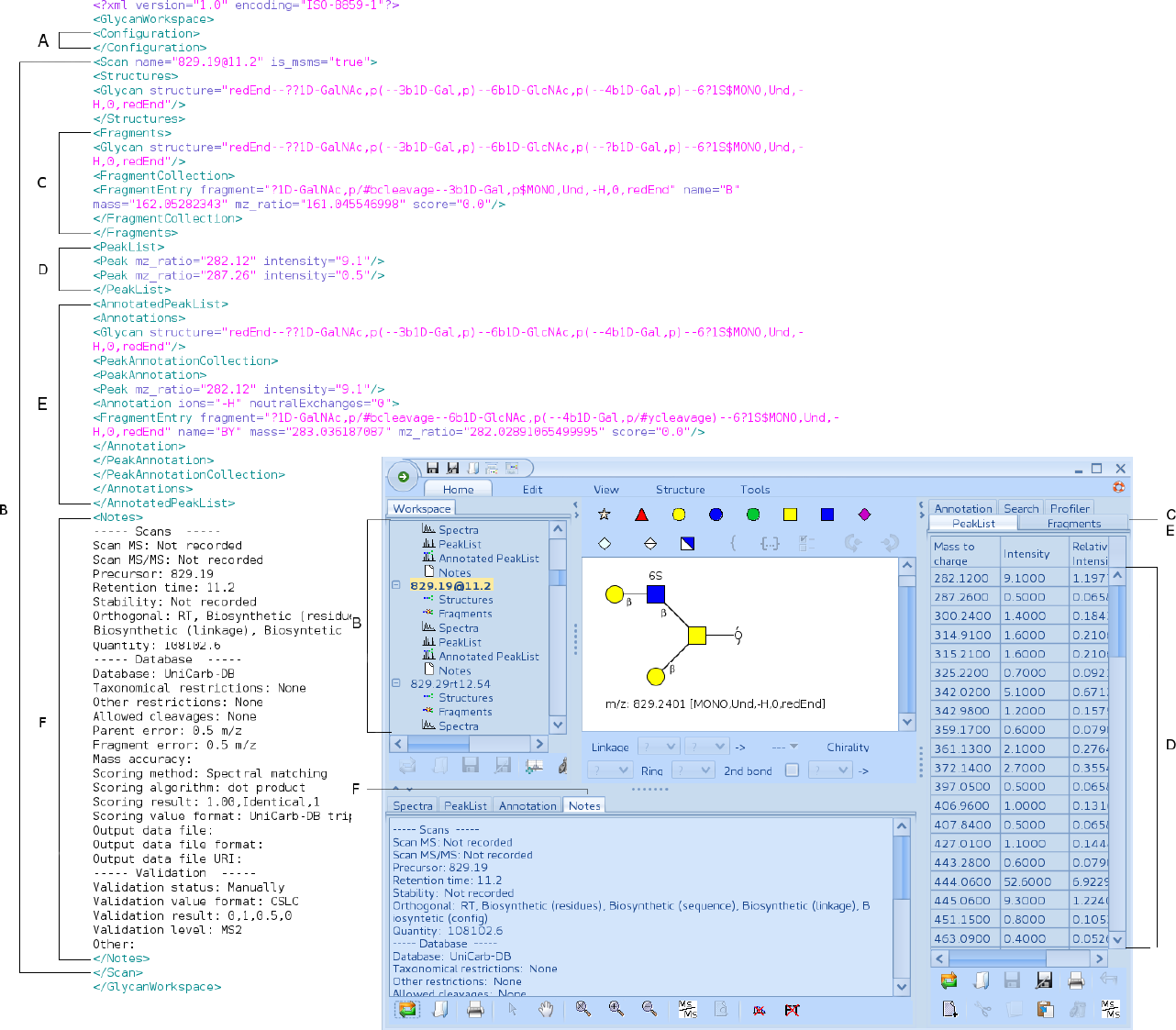
Main sections in the GlycoWorkBench template. The graphical interface of GlycoWorkBench is divided on different panels. The central panel displays the glycan structure in SNGF cartoon notation^41^. Below the structure, multiple panels can be accessed through their respective tab. The last tab corresponds to the ‘Notes’ section (F). Here, additional MIRAGE data can be included. All structures in the file are listed in the pane on the left (B). In this case, the names correspond to their *m/z* values and retention times. The peak list (D) and the list of fragments (C,E) are included on the right side panel. The above mentioned sections are shown as they are stored in the respective .gwp files. GWP files follow XML formatting rules where every element in the file is delimited by tags. The notes for example are defined between the <notes> and </notes> tags (F). Sections can be nested into other sections and be present in multiple instances.

Figure 3 shows the sections that are typically included in a .gwp file. A tag is represented by the “<” and “>”symbols and it defines the different elements in a file. These elements are delimited by a start tag e.g. <scan> and an end tag, e.g. </scan>. The example shown in Figure 3 belongs to a single structure however somewhat simplified, highlighting important MIRAGE tags. In order to be MIRAGE-compatible we introduced a ‘Notes’ section, for recording of orthogonal assignment methods, scoring and validation (see section “*Interpretation of the MIRAGE guidelines for parameter organization”*). The format of the ‘Notes’ section needs to be respected in order to upload its information to UniCarb-DR. The proposed format for the Figure 3 ‘Notes’ section is expanded in Supplementary Material.

### Step 3) MIRAGE record-Native MS files deposit

There is currently no organization comparable to the proteomic PRIDE^36^ and the ProteomeXchange consortium^37^committed to host glycomics/glycoproteomics MS experimental data. Without this option, we conceived a new scheme for how glycomic MS data could be made available. We adapted a web-based proteomic MS Laboratory Information Management System called Proteios Software Environment (http://www.proteios.org)^38^, and using this system, files can be managed and uploaded to storage accessible through WebDAV. The concept implementation is using Swestore (http://www.snic.se/allocations/swestore/) as online storage with regulated file access through dCache (https://www.dcache.org). From Proteios it is subsequently possible to export an entire project with all the required data into appropriate formats for curation. Native MS data (.raw and mzML) is made publicly available at Swestore URIs (Uniform Resource Identifiers) that can be provided in the MIRAGE spreadsheets before submission to UniCarb-DR. Storing of the unprocessed data as part of an open access policy requires long-term commitment from national/international life science data storage organizations. In order to be compliant with MIRAGE guidelines, the organizations or institutions that provide long term data-access that goes beyond individually funded glycomic efforts, will have to be identified. We are currently engaged with JPOST (Japan ProteOme STandard Repository) to develop a pipeline to push unprocessed glycomic MS data from Proteios for permanent storage to a designed repository for glycomic and glycoproteomic data (GlycoPOST, unpublished).

### Step 4) MIRAGE record-Glycan structure registration

GlyTouCan^35^ is a glycan structure repository promoted by the glycocommunity as the prime location for generating unique identifiers for glycan structures. It is recommended that glycan structures should be submitted to this repository as part of a publication of glycomic data. In order to avoid duplicate submissions to both UniCarb-DR and GlyTouCan we developed a tool that assesses whether the structures submitted to UniCarb-DR are already deposited in GlyTouCan. In this case the GlyTouCan ID provides a link to UniCarb-DR. If a UniCarb-DR submitted structure is not available in GlyTouCan, a new ID will be generated and communicated to UniCarb-DR. This process will commence after the submission of data to UniCarb-DR.

### Interpretation of the MIRAGE guidelines for parameter organization

The MIRAGE guidelines are generic and flexible to collect information from different types of experiments aiming to study glycoconjugates. However, the use of commonly defined vocabularies is required for databasing in order to easily compare data within UniCarb-DR as well as shared data with other glycomic and life science databases. To preserve the flexibility of MIRAGE guidelines in the reporting we provide “free text” fields to describe experiments. Only to record key MIRAGE parameters (tissue, MS-device) a rigorous reporting language is implemented. Inspired by the organization of PRIDE^39^, four different types of formats of the MIRAGE parameters were encoded in UniCarb-DR (Table 1) and outlined in Supplementary Material.

**Table 1.** Examples of parameters identified in the MIRAGE guidelines. (1) Defined formats to control the input and presentation of certain data, (2) Ontologies or existing vocabulary that can be adopted from different sources, (3)Parameters that need to be expanded into ontologies or controlled vocabulary to use in glycomics, (4) Free unrestricted text

| **1. Defined formats** |  |
| --- | --- |
| **MIRAGE parameter** | **Format** |
| Unique identifiers for structure and entry | Consecutive number |
| Dates | mm/dd/yyyy |
| Oligosaccharide structure (.gws) | .gws, but also translated into glyco-CT (PMID) and WURCS (PMID) |
| **2. Existing ontologies and controlled vocabulary** |  |
| **Ontologies** | **Origin** |
| Cell lines | http://www.clo-ontology.org/ |
| Species | https://www.ncbi.nlm.nih.gov/taxonomy |
| Tissues | https://meshb.nlm.nih.gov/record/ui?ui=D014024 |
| MS devices | https://www.ebi.ac.uk/ols/ontologies/ms |
| Software | https://www.ebi.ac.uk/ols/ontologies/ms |
| Attached protein-UniProt | http://www.uniprot.org/ |
| **3. Parameters that need to be expanded or created for glycoanalytics** |  |
| **MIRAGE Parameter** | **Partial Information** |
| Treatments and orthogonal methods for isolation/characterisation | Glycosuite (PMID) and Sugarbind (PMID) |
| Software used for data processing and structure assignment |  |
| MS Scoring method |  |
| Validation methods of assigned structure |  |
| **4. Free text** |  |
| Growth/harvest conditions for recombinantly produced material |  |
| Treatments and/or storage conditions for material isolated from tissues |  |
| Synthesis steps for chemically derived material |  |
| Purification steps |  |
| Sprayer features |  |
| Voltages and other parameters relevant for ESI experiments |  |
| Collision energy CID function |  |

### Upload of MIRAGE compatible MS fragmentation spectra to UniCarb-DR

MIRAGE-compliant data sets generated by the web interface along with data stored in .gwp files of both individual fragment spectra and structures can be itself be submitted as supplementary data in a given publication. We also propose uploading theses collected and structured glycomic information (spreadsheets and .gwp files). The upload allows fully and partly assigned structures as can be seen in Figure 4. The reporting of orthogonal methods (i.e. NMR, HPLC retention time mapping, and chemical/enzymatic treatment) also justifies that UniCarb-DR accepts structures, where MS but not MS^n^ has been collected. In Figure 4 there are examples from reference 1 in UniCarb-DR (http://unicarb-dr.biomedicine.gu.se/references/1) of assigned structures both with associated fragment data and structures without fragment data. The latter structures would have been assigned based on retention time (RT) and biosynthetic knowledge about the constituting monosaccharides, linkage position and configuration. We have assembled an expandable list of treatments and orthogonal methods for isolation/characterisation (Table 1 and supplementary spreadsheet). Current records in UniCarb-DR have been uploaded using data generated in various laboratories by researchers in the author list. During this process we noted that the MIRAGE guidelines were focused on the overall description of the experiment to encourage the use of the submission tool implementing the MIRAGE guidelines. However, the requirement to record the full information about individual structures (e.g. scoring and orthogonal method validation) is time consuming. Hence, UniCarb-DR is also accepting data with only partial MIRAGE records for an individual structure, i e. at least the record of the parent ion mass has to be provided, excluding information about scoring and validation.

**Figure 4.**
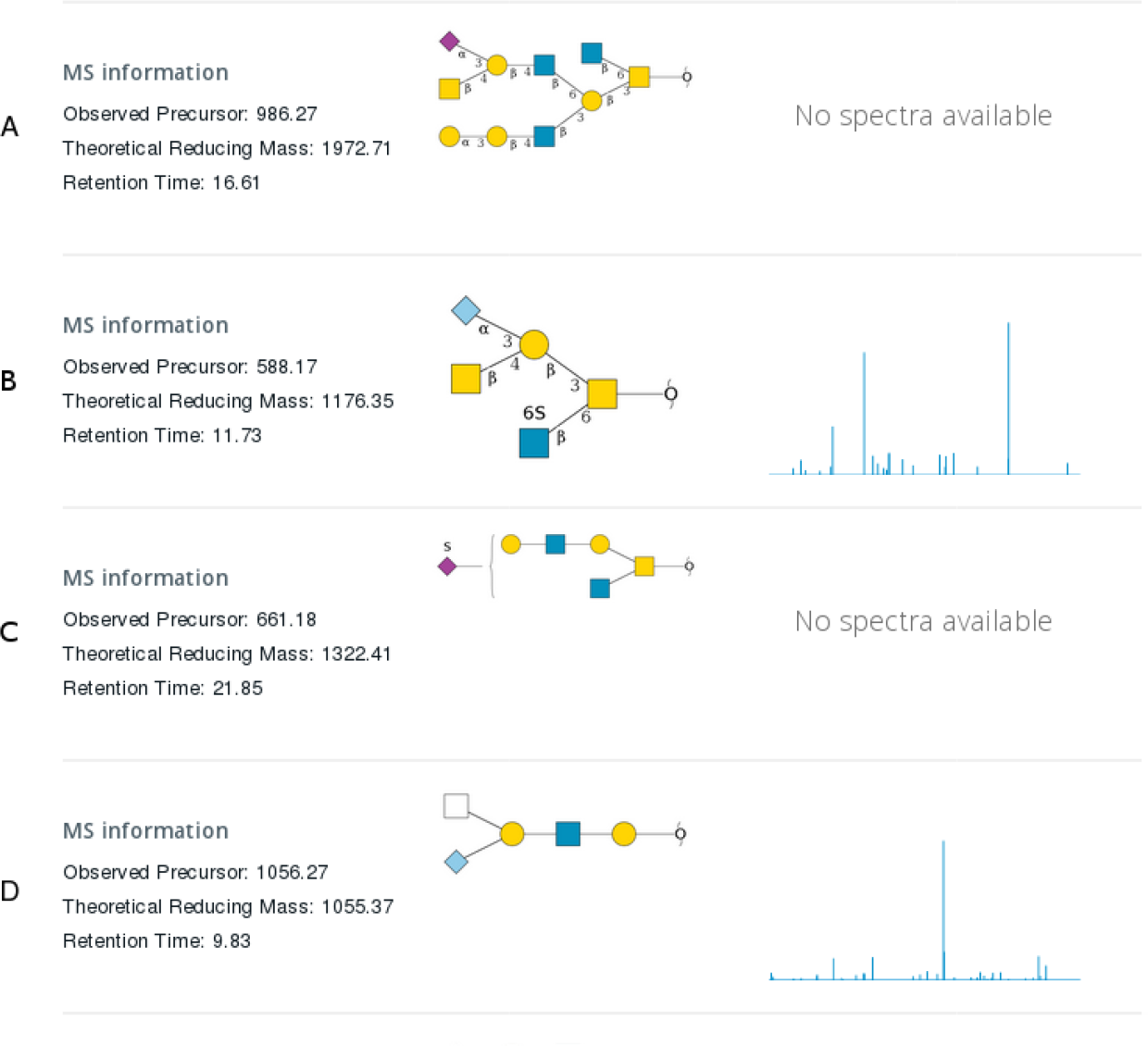
Different types of entries managed by the UniCarb-DR submission tool. A) A structure proposed from LC-MS without LC-MS^2^ spectrum. B) Fully assigned structure with associated LC-MS^2^ spectrum. C) Partially assigned structure proposed from LC-MS without LC-MS^2^ spectrum. D) Partially assigned structure with associated LC-MS^2^ spectrum.

## DISCUSSION

The lack of an established formalized description of glycomic experiments may cause progress in the field to stall. The community needs to rely on sharing yet it remains one open question as to how? Here, we are proposing a solution for a glycomics MS format using spreadsheets in combination with GlycoWorkbench files. This proposed format is one step closer to enforcing MIRAGE compliant scientific publications in glycomics. Past experience in introducing guidelines for glycomic studies as part of publications (http://www.mcponline.org/site/misc/glycomic.xhtml) has shown that if there is a clear pathway and format, researchers will conform. With the developed tools in this report, we propose that glycomic MS recording can be adopted early at the start of a project, where the spreadsheet can be completed and modified as the project evolves. Furthermore, the use of GlycoWorkbench files for saving glycomic structural discoveries can be implemented for housekeeping. The spreadsheet and .gwp format are flexible to support a variety of glycomics MS applications only with limited modifications using the templates provided. Hence, journal editors can confidently request that authors are MIRAGE compatible by providing the files as supplementary data. With an increasing awareness of MIRAGE formats, a grassroots movement from those performing both research and scientific publication reviewing will insist that, not only their own but also others, data are MIRAGE compliant in scientific publications.

While both spreadsheets and .gwp formats are flexible and can easily be expanded to adapt to various workflows as part of a publication supplement, the UniCarb-DR upload requires a known number and defined formats for the variables. Hence, an e-upload will have to be further developed to support various glycomic workflows, in addition to the current LC-MS and MS module. The GlycoWorkbench structure format has been adopted in other glycomic commercial (Glycoquest, Bruker, Bremen Germany) and academic (GRITS Toolbox (http://www.grits-toolbox.org/) software projects. Hence, automated submission to UniCarb-DR is likely to be easily implemented for these. Other workflows utilized in glycomics, such as permethylation followed by MS with or without coupled separation, are also easily implemented if the spectra are from single isomers. With several isomers present in one spectrum, these data can still be recorded in GlycoWorkbench (several structures recorded in one “Scan”), but to accommodate upload, the UniCarb-DR format will need to be modified. Similar concerns relate to workflows involving multiple MS^n^, even though the flexibility of GlycoWorkbench allows this to be recorded as sub-“Scans”. Other glycomic MS workflow (eg ion mobility MS) may need smaller or larger adoption of UniCarb-DR. We request the help from the community to identify additional major glycomic workflows for us to adapt the submission accordingly. Templates for these alternative workflows may require modifications regarding software and experiment. We are also planning to modify the Glycoforest^32^ output to allow automated submission to UniCarb-DR. The Glycoforest module generating consensus spectra, could be used in the curation process of UniCarb-DR. Furthermore, the Glycoforest module for matching and clustering spectra could use the repository and score similarities between samples. This could be achieved through access to stored native MS data from an outside location with associated metadata.

The commitment to store glycomic MS datasets is essential. It is obvious that interpretation of glycomic LC-MS data is based only on the knowledge of the interpreters^32^, be it a software or a human researcher or both. Hence, glycomic native MS data should be considered as libraries that could be read repeatedly to harvest new data and to ask new questions. This is even more important when glycomics evolves similarly to proteomics, where data independent acquisition^40^ could be utilized as a means to generate data from clinical or other reference samples. These glycomic libraries could be used for harvesting information for hypothesis driven glycomics. Similar to the PRIDE and ProteomeXchange initiatives, the glycomic community needs to voice the unanimous opinion that this is needed and target both national and international life science e-infrastructure organizations. The MIRAGE board already identified this requirement by introducing the demand for raw data deposition in the guidelines.

A pipeline for curation of data between UniCarb-DR to UniCarb-DB is changing the perception of how curated glycomic structural databases will be generated. The top-down approach of a database generator and curator trolling literature for information will shift to researchers submitting and managing their own data. Researchers and curators will need software tools to help in the curation process. Utilization of the metadata in the reporting guidelines together with evaluation of accompanying publication as well as previous and future knowledge can together aid the curation process. This process must remain objective and transparent in that information can only be added neither deleted nor altered without permission from the data supplier. The curation process will be strengthened by the MIRAGE information in order to generate unbiased statement about data quality. The mission of UniCarb-DR and -DB is to support the development of a knowledgebase of glycan structures by providing the pipeline for glycomic experimental MS data.

## Acknowledgements

This work was financed by the European Union FP7 GastricGlycoExplorer ITN (No 316929), the Swedish Research Council (621-2013-5895), The Swedish Foundation for International Cooperation in Research and Higher Education (STINT) initiation grant (IB2015-5931) and institutional grant (IG2010-2050). SIB is supported by the Swiss Federal Government through the State Secretariat for Education, Research and Innovation (SERI). ExPASy is maintained by the web team of the Swiss Institute of Bioinformatics and hosted at the Vital-IT Competency Center. The MIRAGE project is supported by Beilstein-Institut.

Author contributions: M.A.R.-M. designed and lead the development of the database; J.M. O.H, and P.A. contributed to the UniCarb-DR implementation and designed the database and web interface; C.J. and N.G.K. contributed to the design of the upload forms, C.J., V.V., K.M., C.A., R.L.M, T.Z. contributed with data for beta-testing of the data submission tool and feedback; N.P.A, D.S. and K.F.A.-K. contributed to the setting up the communication with GlyTouCan; F.Le. contributed with design of Proteios for glyco MS data; W.B.S, D.K., P.M.R., M.W, C.K., N.H.P, K.F.A.-K, F. Li. and N.G.K. formed the advisory core that contributed with feedback on repository design, advised in implementation of glycomic MS/reporting guidelines and design of databases and coding; K.F.A.-K and N.G.K jointly received the funds for UniCarb-DR and GlyTouCan communication; F.Li. and N.G.K. jointly received funds for development of the UniCarb-DR infrastructure and structured the manuscript; N.G.K. proposed the UniCarb-DR concept and managed the project. M.A.R.-M. F.Li and N.G.K. jointly wrote the manuscript with input from the other authors. All authors signed of on the final version.

## References

1. Aebersold R, et al. How many human proteoforms are there? Nat Chem Biol 14, 206–214 (2018).

2. Varki A. Glycan-based interactions involving vertebrate sialic-acid-recognizing proteins. Nature 446, 1023–1029 (2007).

3. Rudd PM, Elliott T, Cresswell P, Wilson IA, Dwek RA. Glycosylation and the immune system. Science 291, 2370–2376 (2001).

4. Sparks SE, Krasnewich DM. Congenital Disorders of N-Linked Glycosylation and Multiple Pathway Overview. In: GeneReviews((R)) (ed^(eds Adam MP, et al.) (1993).

5. Jaeken J, Matthijs G. Congenital disorders of glycosylation. Annu Rev Genomics Hum Genet 2, 129–151 (2001).

6. Wuhrer M. Glycomics using mass spectrometry. Glycoconj J 30, 11–22 (2013).

7. Zaia J. Mass spectrometry and the emerging field of glycomics. Chem Biol 15, 881–892 (2008).

8. An HJ, Kronewitter SR, de Leoz ML, Lebrilla CB. Glycomics and disease markers. Curr Opin Chem Biol 13, 601–607 (2009).

9. Brazma A, et al. Minimum information about a microarray experiment (MIAME)-toward standards for microarray data. Nat Genet 29, 365–371 (2001).

10. Taylor CF, et al. The minimum information about a proteomics experiment (MIAPE). Nat Biotechnol 25, 887–893 (2007).

11. Tipton KF, et al. Standards for Reporting Enzyme Data: The STRENDA Consortium: What it aims to do and why it should be helpful. Perspectives in Science 1, 131–137 (2014).

12. Jenkins H, et al. A proposed framework for the description of plant metabolomics experiments and their results. Nat Biotechnol 22, 1601–1606 (2004).

13. Goodacre R, et al. Proposed minimum reporting standards for data analysis in metabolomics. Metabolomics 3, 231–241 (2007).

14. McQuilton P, et al. BioSharing: curated and crowd-sourced metadata standards, databases and data policies in the life sciences. Database 2016, baw075–baw075 (2016).

15. York WS, et al. MIRAGE: the minimum information required for a glycomics experiment. Glycobiology 24, 402–406 (2014).

16. Struwe WB, et al. The minimum information required for a glycomics experiment (MIRAGE) project: sample preparation guidelines for reliable reporting of glycomics datasets. Glycobiology 26, 907–910 (2016).

17. Kolarich D, et al. The minimum information required for a glycomics experiment (MIRAGE) project: improving the standards for reporting mass-spectrometry-based glycoanalytic data. Mol Cell Proteomics 12, 991–995 (2013).

18. Liu Y, et al. The minimum information required for a glycomics experiment (MIRAGE) project: improving the standards for reporting glycan microarray-based data. Glycobiology, (2016).

19. Packer NH, et al. Frontiers in glycomics: bioinformatics and biomarkers in disease. An NIH white paper prepared from discussions by the focus groups at a workshop on the NIH campus, Bethesda MD (September 11-13, 2006). Proteomics 8, 8–20 (2008).

20. Doubet S, Bock K, Smith D, Darvill A, Albersheim P. The Complex Carbohydrate Structure Database. Trends Biochem Sci 14, 475–477 (1989).

21. Ranzinger R, Herget S, Wetter T, von der Lieth CW. GlycomeDB - integration of open-access carbohydrate structure databases. BMC Bioinformatics 9, 384 (2008).

22. Lutteke T, Bohne-Lang A, Loss A, Goetz T, Frank M, von der Lieth CW. GLYCOSCIENCES.de: an Internet portal to support glycomics and glycobiology research. Glycobiology 16, 71R–81R (2006).

23. Cooper CA, Harrison MJ, Wilkins MR, Packer NH. GlycoSuiteDB: a new curated relational database of glycoprotein glycan structures and their biological sources. Nucleic Acids Res 29, 332–335 (2001).

24. Campbell MP, et al. UniCarbKB: putting the pieces together for glycomics research. Proteomics 11, 4117–4121 (2011).

25. Hayes CA, et al. UniCarb-DB: a database resource for glycomic discovery. Bioinformatics 27, 1343–1344 (2011).

26. Campbell MP, et al. Validation of the curation pipeline of UniCarb-DB: building a global glycan reference MS/MS repository. Biochim Biophys Acta 1844, 108–116 (2014).

27. Joshi HJ, Harrison MJ, Schulz BL, Cooper CA, Packer NH, Karlsson NG. Development of a mass fingerprinting tool for automated interpretation of oligosaccharide fragmentation data. Proteomics 4, 1650–1664 (2004).

28. Apte A, Meitei NS. Bioinformatics in glycomics: glycan characterization with mass spectrometric data using SimGlycan. Methods Mol Biol 600, 269–281 (2010).

29. He L, Xin L, Shan B, Lajoie GA, Ma B. GlycoMaster DB: software to assist the automated identification of N-linked glycopeptides by tandem mass spectrometry. J Proteome Res 13, 3881–3895 (2014).

30. Ashline DJ, Hanneman AJ, Zhang H, Reinhold VN. Structural documentation of glycan epitopes: sequential mass spectrometry and spectral matching. J Am Soc Mass Spectrom 25, 444–453 (2014).

31. Sun W, Lajoie GA, Ma B, Zhang K. A Novel Algorithm for Glycan de novo Sequencing Using Tandem Mass Spectrometry. (ed^(eds). Springer International Publishing (2015).

32. Horlacher O, et al. Glycoforest 1.0. Anal Chem, (2017).

33. Jansen BC, et al. MassyTools: A High-Throughput Targeted Data Processing Tool for Relative Quantitation and Quality Control Developed for Glycomic and Glycoproteomic MALDI-MS. J Proteome Res 14, 5088–5098 (2015).

34. Damerell D, Ceroni A, Maass K, Ranzinger R, Dell A, Haslam SM. The GlycanBuilder and GlycoWorkbench glycoinformatics tools: updates and new developments. Biol Chem 393, 1357–1362 (2012).

35. Tiemeyer M, et al. GlyTouCan: an accessible glycan structure repository. Glycobiology 27, 915–919 (2017).

36. Vizcaino JA, et al. 2016 update of the PRIDE database and its related tools. Nucleic Acids Res 44, D447–456 (2016).

37. Vizcaino JA, et al. ProteomeXchange provides globally coordinated proteomics data submission and dissemination. Nat Biotechnol 32, 223–226 (2014).

38. Hakkinen J, Vincic G, Mansson O, Warell K, Levander F. The proteios software environment: an extensible multiuser platform for management and analysis of proteomics data. J Proteome Res 8, 3037–3043 (2009).

39. Jones P, et al. PRIDE: a public repository of protein and peptide identifications for the proteomics community. Nucleic Acids Res 34, D659–663 (2006).

40. Gillet LC, Leitner A, Aebersold R. Mass Spectrometry Applied to Bottom-Up Proteomics: Entering the High-Throughput Era for Hypothesis Testing. Annu Rev Anal Chem (Palo Alto Calif) 9, 449–472 (2016).

41. Varki A, et al. Symbol Nomenclature for Graphical Representations of Glycans. Glycobiology 25, 1323–1324 (2015).

